# Power analysis of artificial selection experiments using efficient whole genome simulation of quantitative traits

**DOI:** 10.1101/005892

**Authors:** Darren Kessner, John Novembre

## Abstract

Evolve and resequence studies combine artificial selection experiments with massively parallel sequencing technology to study the genetic basis for complex traits. In these experiments, individuals are selected for extreme values of a trait, causing alleles at quantitative trait loci (QTLs) to increase or decrease in frequency in the experimental population. We present a new analysis of the power of artificial selection experiments to detect and localize quantitative trait loci. This analysis uses a simulation framework that explicitly models whole genomes of individuals, quantitative traits, and selection based on individual trait values. We find that explicitly modeling QTL provides produces qualitatively different insights than considering independent loci with constant selection coefficients. Specifically, we observe how interference between QTLs under selection impacts the trajectories and lengthens the fixation times of selected alleles. We also show that a substantial portion of the genetic variance of the trait (50–100%) can be explained by detected QTLs in as little as 20 generations of selection, depending on the trait architecture and experimental design. Furthermore, we show that power depends crucially on the opportunity for recombination during the experiment. Finally, we show that an increase in power is obtained by leveraging founder haplotype information to obtain allele frequency estimates.

## Introduction

Some of the first artificial selection experiments, performed in the early 1900s, were designed to settle ongoing debates about the nature of selection. In particular, early researchers hoped to answer questions about whether selection on continuous variation was even possible, and how to reconcile this with the Mendelian viewpoint of genes as discrete heritable units (see Falconer (1992) for a history of early selection experiments, and Garland and Rose (2009) for a comprehensive overview of experimental evolution).

More recently, researchers have used artificial selection experiments to study the genetic basis for complex traits by analyzing allele frequency changes in evolved populations. This technique has been used in a wide variety of organisms, including for example maize (Laurie *et al.*, 2004), yeast (Ehrenreich *et al.*, 2010), fruit flies (Nuzhdin *et al.*, 2007; Teotonio *et al.*, 2009), chickens (Johansson *et al.*, 2010), and mice (Keightley and Bulfield, 1993).

Massively parallel (also known as next-generation) sequencing of pooled samples has enabled researchers to obtain genome-wide allele frequency estimates for a population in a cost-effective manner (Futschik and Schlotterer, 2010). These technological advances have led to the development of the Evolve and Resequence (E&R) method for mapping traits (Burke *et al.*, 2010; Turner *et al.*, 2011; Parts *et al.*, 2011; Orozco-terWengel *et al.*, 2012; Remolina *et al.*, 2012). In E&R studies, artificial selection is followed by pooled sequencing of genomic DNA from multiple individuals. A site exhibiting a large allele frequency change in the selected population suggests the presence of a nearby quantitative trait locus (QTL) (Figure 1).

**Figure 1.**
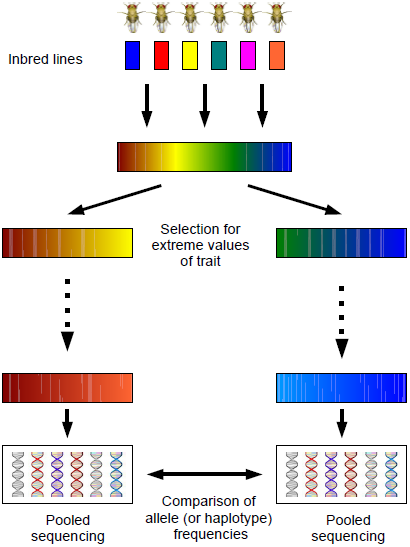
Evolve and resequence experiment. After initial neutral mixing of founders, individuals with extreme values of the trait are selected to create the next generation. After several generations of selection, populations are sequenced and analyzed for allele frequency differences.

Due to limitations on resources and time, researchers will face many design decisions as they set up artificial selection experiments. First, the size of experimental populations affects the degree of genetic drift and the efficacy of selection. Next, the strength of selection is determined by the proportion of individuals selected each generation to create the next generation. In addition, the length of the experiment must be chosen in conjunction with the strength of selection so as to maximize allele frequency differences at QTLs, while at the same time minimizing differences at neutral loci due to drift. For example, extremely strong selection over a large number of generations will result in the fixation of variants across the genome, which will stifle any ability to distinguish QTLs from neutral loci. An additional factor consider is that in the selection can be performed on a single population which will be compared to a control population; alternatively, selection can be divergent, where two populations selected in opposite directions will be compared. Finally, replication is a key consideration in any experiment, and especially in experimental evolution, where random genetic drift plays a large role: observation of a large allele frequency difference at a locus in multiple replicate experiments will increase confidence that the difference is due to selection at the locus rather than drift.

Previous studies of the power of artificial selection experiments to detect trait loci have taken the traditional population genetics approach in which selection is parametrized using selection coefficients that remain constant each generation. For example, previous work by Kim and Stephan (1999) analyzed a single locus under a constant selection coefficient using a diffusion approximation, and two recent simulation studies employed forward simulations with loci parametrized by constant selection coefficients to obtain power estimates (Kofler and Schlotterer, 2014; Baldwin-Brown *et al.*, 2014).

However, E&R experiments lie at the interface between population genetics and quantitative genetics. The selection pressure on a QTL is more naturally parametrized using concepts from quantitative genetics such as effect sizes, genetic variance, and heritability, in addition to experimental design parameters such as the proportion of individuals selected each generation. Furthermore, as we will see, the effect of a QTL allele on an individual’s fitness depends strongly on interaction with the other QTL alleles carried by the individual. As noted by Felsenstein (1974, 1987), finite populations necessarily experience random linkage disequilibrium between any two polymorphic loci. This linkage disequilibrium results in interference, where the effect of selection on each locus is decreased (Hill and Robertson, 1966).

In this study, we introduce a new simulation framework to investigate the power of artificial selection experiments to detect and localize QTLs contributing to a quantitative trait. Our simulations employ a whole-genome quantitative genetic model of loci underlying a quantitative trait, and we explicitly model artificial selection of individuals each generation based on trait values. As in Kofler and Schlotterer (2014) and Baldwin-Brown *et al.* (2014), we model *Drosophila melanogaster* individuals, and we use experimental parameters similar to those used by Turner and Miller (2012).

Our simulations show that forward simulations of a locus assuming a constant selection coefficient do not fully capture the allele frequency dynamics of a QTL under artificial selection on a quantitative trait. In contrast, explicit quantitative genetic modeling of the trait leads to insights regarding the effect of the trait architecture on the allele frequency trajectory of a QTL. For instance, by simulating the entire genome of individuals, we demonstrate the important role that recombination plays in decreasing interference between QTLs, as well as reducing linkage disequilibrium between QTLs and neighboring neutral loci. Our results emphasize that designing the artificial selection experiment to allow more opportunity for recombination increases the ability to detect and localize QTLs.

Finally, previous work suggested that when founder sequence information is available, one can obtain more accurate allele frequency estimates by estimating local haplotype frequencies from pooled read data (Kessner *et al.*, 2013). We show that these improved allele frequency estimates, when compared to estimates calculated directly from read counts, lead to an increase in power.

## Results

### Comparison between selection coefficient simulations and explicit quantitative genetic modeling

To address the importance of modeling quantitative traits explicitly, we first investigated allele frequency trajectories of a focal QTL contributing to a quantitative trait under truncation selection in comparison to a single locus with 2 alleles with a constant additive selection coefficient.

We simulated populations of size *N* = 1000 for the selection coefficient simulations. For the artificial selection simulations, we simulated populations of size *N* = 5000 with the top 20% of individuals selected based on their trait vales to create the next generation, so that the effective population size was *N_e_* = 1000. We ran the artificial selection simulations with 12 canonical trait architectures with various heritabilities and QTL counts, and with exponentially distributed effect sizes (see Methods). In all simulations, the initial frequency of trait-increasing allele was .2, and selection is assumed to be in the direction of increasing trait values.

As seen in Figure 2 (A and C), the allele frequency trajectories of the focal QTL under truncation selection are qualitatively different from the trajectories under a constant selection coefficient. While a strong selection coefficient leads to nearly deterministic allele frequency trajectories, trajectories of a focal QTL under strong truncation selection are dependent on the underlying trait architecture. In particular, the focal QTL may experience interference from other linked QTLs. This effect can be seen in the trajectories where the focal QTL decreases in frequency at first, due to repulsion linkage disequilibrium (i.e. QTL of opposite effect are positively associated), but eventually increases in frequency once linkage disequilibrium has decreased sufficiently through recombination.

**Figure 2.**
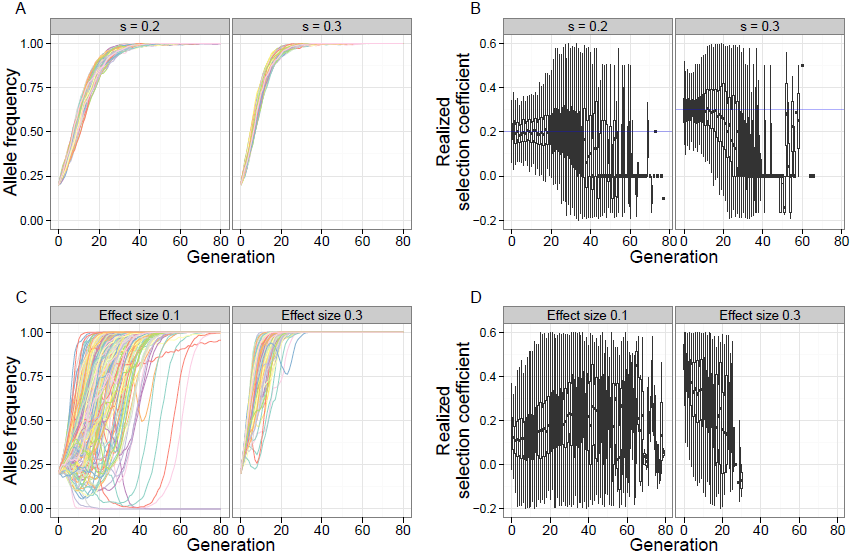
Qualitative differences between fixed selection coefficient and truncation selection on a quantitative trait. A focal QTL exhibits fundamentally different behavior under a constant selection coefficient, compared to truncation selection on a quantitative trait. Shown are allele frequency trajectories (A) and realized selection coefficient distributions (B) of a locus under two different selection coefficients (.2, .3) [N=1000, 80 generations]. C and D show the same for a focal QTL (effect sizes .1, .3) under truncation selection on the trait [N=5000, 20% selected, 80 generations, 100 QTLs, *h*^2^ = .8, *σ*^2^*_trait_* = 1].

In addition, once an allele with a constant selection coefficient reaches high frequency, it only gradually increases in the final generations before finally going to fixation. In contrast, the focal QTL under truncation selection tended to become fixed quickly after reaching high frequency in the population. This behavior is not surprising, because after a few generations of selection, the upper tail of the population trait value distribution is highly enriched for individuals carrying high-effect variants.

To further illustrate these qualitative differences, we analyzed the realized selection coefficient of the trajectories, which represents the selection coefficient that would result in a given single-generation allele frequency change under a deterministic model (see Methods). Under a constant selection coefficient, the mean realized selection coefficient tracks the true selection coefficient closely during the selection phase, after which it decreases to zero during the drift phase (Figure 2B). Under truncation selection, the behavior of the mean realized selection coefficient depends on the underlying genetic architecture of the trait. When the effect size is low, the realized selection coefficient increases each generation – this is because selection acts on the larger effect QTLs first, and then has a greater effect on the focal QTL after the larger effect QTLs have reached fixation. On the other hand, when the effect size is higher, the focal QTL experiences very strong selection initially, decreasing as the focal QTL rises to fixation (Figure 2D).

Another effect of the underlying trait architecture can be seen in the fixation times of the focal QTL (Figure 3). For a given effect size and heritability, the fixation time of the focal QTL increases with total number of QTLs due to interference.

**Figure 3.**
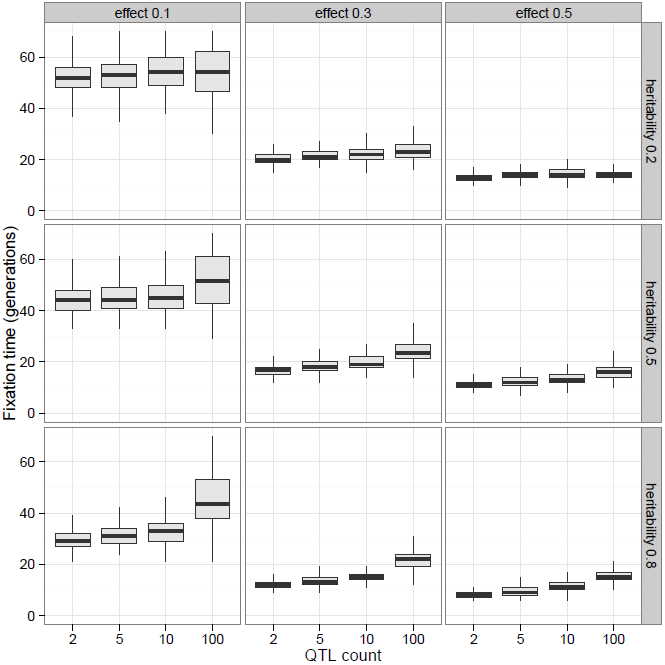
Effect of genetic architecture on fixation times. Fixation times of a focal QTL for a trait under truncation selection decrease with increasing effect size and heritability. For a fixed effect size and heritability, the fixation times increase with the number of QTLs contributing to the trait due to interference. [N=5000, 20% selected, 80 generations]

### Measurement of power to detect and localize QTLs

In all of the following analyses, we examine the power to detect QTLs through allele frequency differences at variant sites. There is currently no consensus regarding the choice of statistic in the analysis of artificial selection experiments. Both for simplicity and becuase of its use in practice (Parts *et al.*, 2011; Turner and Miller, 2012) we chose to base our analysis on the absolute allele frequency difference *D* = |*p*_1_ − *p*_2_|, where *p*_1_ and *p*_2_ are the allele frequencies of the variant in populations 1 and 2, respectively. We also note that this statistic, also called the difference in derived allele frequencies (Δ*DAF* or *DDAF*), is used in several related or composite methods for detecting selection (Turner *et al.*, 2011; Grossman *et al.*, 2010; Utsunomiya *et al.*, 2013).

We calculate *D* for each variant site in the genome, and we call a site detected if the *D* value exceeds a threshold value. By varying the threshold, we obtain receiver operating characteristic (ROC) curves showing the relationship between power (true positive rate) and the false positive rate.

Due to linkage and strong selection, detected QTLs will generally have neighboring neutral variants whose allele frequency differences also exceed the detection threshold. Because of this, detection and localization of a QTL are necessarily intertwined. In an actual experimental setting, the entire genomic region surrounding the significantly diverged loci would often be chosen for follow-up studies.

We explored several methods for calculating and interpreting power and false positive rate. We present our results using the method that we found to be most interpretable, and which we summarize here (see Methods for full details on the different methods). For a given *D* value threshold, we determine a *detection region* that consists of all variants within a specified radius of any variant above the threshold (10kb for the results presented here; see Figure 9 for an illustration). Power is calculated as the proportion of genetic variance (in the founder population) expained by the QTLs located within the detection region. We define the *true region* to consist of all variants within the specified radius of any QTL (i.e. 10kb here), and we define the *neutral genome* to be the rest of the genome (outside the true region). False positive rate is calculated as the proportion of the neutral genome covered by the detection region. Thus, a false positive rate of .01 can be interpreted to mean that 1% of the genome would be incorrectly flagged for follow-up studies. In *Drosophila melanogaster*, this would correspond to a 1.4mb region.

### Advantages of divergent artificial selection

In artificial selection studies of quantitative traits, one commonly used technique is divergent (also called bidirectional) selection, where the first “high” population is obtained by selecting from the upper tail of the trait value distribution each generation, and the second “low” population is obtained by selecting from the lower tail (for example Johansson *et al.* (2010) and Turner and Miller (2012)). Use of this technique presumes that allele frequency differences between the high and low lines will be more pronounced at QTLs contributing to the trait than, for example, differences between the high line and a control population that has been evolving neutrally.

To compare the power obtained by a divergent selection experiment to the power obtained by selection in a single direction, we simulated 3 populations originating from a single founder population, where one population was selected for high values, one selected for low values, and one allowed to evolve neutrally. Each population had 1000 individuals, of which 20% were selected each generation in the high and low populations, for 20 generations total.

We ran the simulations for each of our 12 canonical architectures (see Methods), and we show the results in Figure 4 for some of these (*h*^2^ = .5, with 5, 10, and 100 QTLs). Comparison between the high and low populations leads to a substantial increase in power over the comparison between the high and neutral populations. In particular, at a 1% false positive rate, roughly half of the genetic variance of the trait is detected in the high vs. low comparison but not in the high vs. neutral comparison.

**Figure 4.**
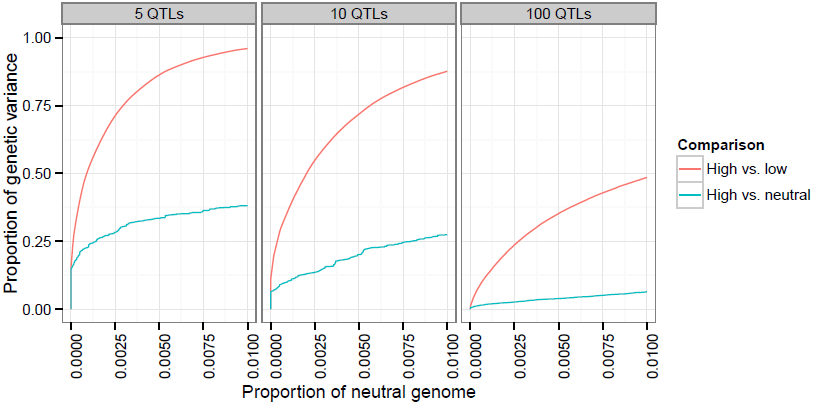
Increase in power due to divergent selection. Comparison of two populations divergently selected for extreme values of a trait has greater power to detect QTLs than comparison between selected and control populations. Three populations of 1000 individuals (high, low, neutral) originating from a single founder population were simulated for 20 generations, with 20% selected each generation in the high and low populations. Shown are the scenarios with 5, 10, and 100 QTLs, with *h*^2^ = .5.

### Extent of increased power due to replication

Another technique available in artificial selection experiments is the use of replicate high and low populations to increase confidence that allele frequency differences between diverged populations are due to selection rather than genetic drift. A natural generalization of the allele frequency difference *D* for replicate populations is 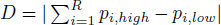, where *R* is the number of replicate population pairs.

We investigated the effects of evolving replicate populations by simulating 5 pairs of populations originating from a single founder population [10 QTLs, *h*^2^ = .5, N=1000, 20% selected, 20 generations].

We then calculated the average power to detect QTLs using subsets of the data representing population replicates from 1 to 5 pairs.

We found that using 2 replicate populations substantially increases power to detect QTLs (Figure 5). For example, at the low false positive rate (neutral genome proportion) of .0005, the proportion of genetic variance detected increases from 25% to 50%. Adding further replicate populations continues to increase power, but with diminishing returns.

**Figure 5.**
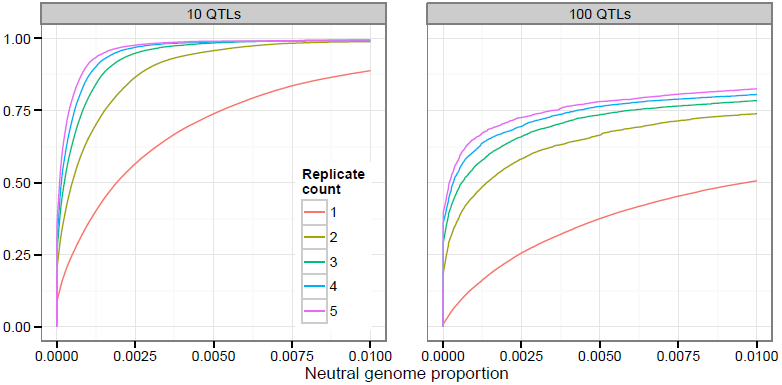
Increase in power due to replicate populations. Adding replicate pairs of divergently selected populations increases power to detect QTLs. 5 pairs of populations originating from a single founder population were simulated [*h*^2^ = .5, N=1000, 20% selected, 20 generations].

### Effect of population size

It is well known that selection on a single locus, defined by a selection coefficient *s*, acts more efficiently in larger populations, as can be seen in the dependence of fixation probabilities and fixation times on the population-scaled selection coefficient 4*N s* (Ewens, 2004).

To investigate how the population size affects artificial selection experiments, we performed simulations of populations of different sizes (*N* = 500, 1000, 2000, 5000) under identical experimental conditions (20% selected, 20 generations), with each of our 12 canonical trait architectures.

In all cases we found a substantial increase in power to detect and localize QTLs as we increased the population size. For example, in the case of 10 QTLs and *h*^2^ = .5, increasing the population size from 500 to 1000 resulted in an additional 25% of the genetic variance detected at the .5% false positive rate (Figure 6). Moreover, while less than 50% of the genetic variance was detected with *N* = 500 at that false positive rate, nearly 100% was detected with *N* = 5000.

**Figure 6.**
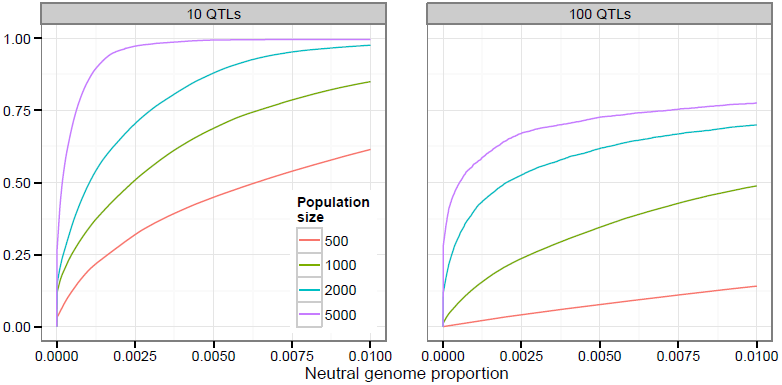
Increase in power due to population size. Using larger population sizes substantially increases power to detect QTLs [*h*^2^ = .5, 20% selected, 20 generations].

### Effects of the length of experiment and proportion selected

We next investigated the effects of the length of the experiment and the strength of selection on the power to detect QTLs. To do this, we simulated 4 different selection scenarios, from weak selection (top/bottom 80% selected each generation) to strong selection (20% selected). Each simulation ran for 80 generations, and we examined snapshots of each population at generations 20, 40, 60, and 80. In these simulations, the populations consisted of 1000 individuals, and the trait had 10 QTLs, with a heritability of .5.

One particularly interesting finding was that under strong selection, power can actually decrease with the number of generations, depending on the false-positive rate threshold chosen (for example, 20% or 40% selected, at false-positive rate .01 in Figure 7). In these scenarios, most of the QTLs have differentiated by generation 20 (over 75% of the genetic variance was detected at false positive rate 1%). However, the lower effective population size induced by strong selection results in the fixation of many neutral variants, which are then falsely detected as QTLs.

**Figure 7.**
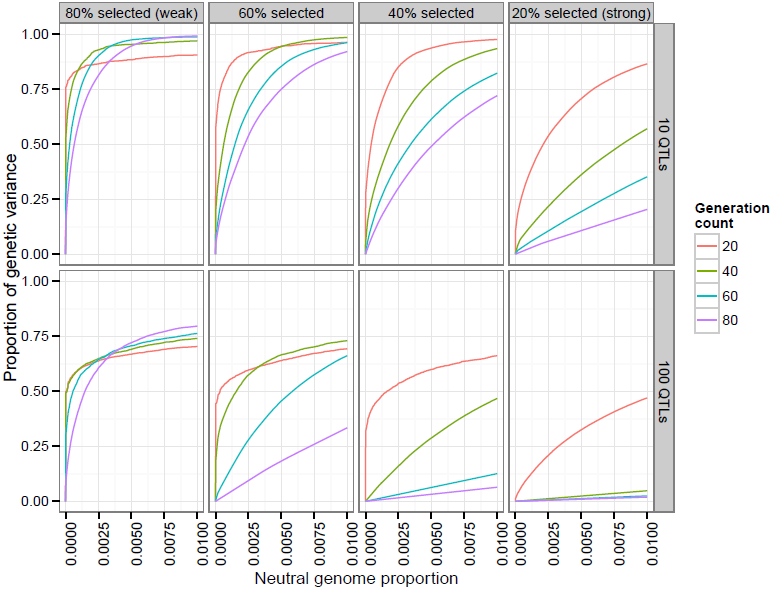
Effects of length of experiment and selection strength. Experiments with weak selection (top/bottom 80% selected each generation) over a longer number of generations have higher power than those with strong selection (20% selected). [N=1000, *h*^2^ = .5]

Conversely, by lowering the selection pressure (80% selected each generation) the maximum power is actually increased. In addition, letting the experiment run for a greater number of generations at this lower selection pressure increases the maximum power attained.

These observations suggest that recombination plays a large role in the power to detect and localize QTLs: reducing the selection pressure and increasing the number of generations should allow more recombination, which reduces linkage disequilibrium between QTLs and neighboring neutral variants. Increased recombination will also reduce interference between QTLs, which should allow lower effect QTLs to be selected and detected. This led us to investigate the effects of recombination further in our subsequent analyses.

### Effect of recombination

To investigate the effects of recombination on the power to detect and localize QTLs, we first considered the effect of increasing the recombination rate of the simulated individuals. We simulated populations of 1000 individuals, with each our 12 canonical trait architectures, with recombination rates at 1x, 2x, 3x, and 4x our standard recombination rate (see Methods) [20 generations of neutral mixing, 20 generations of selection, 20% selected].

We found that increasing the recombination rate does indeed increase the power to distinguish QTLs from neutral variants, for all trait architectures (Figure 8AD). While in practice it is not feasible to experimentally increase the recombination rate of individuals, the experiment can be designed so as to increase the opportunity for individual chromosomes to recombine and thus decrease linkage between QTLs and neighboring neutral variants.

**Figure 8.**
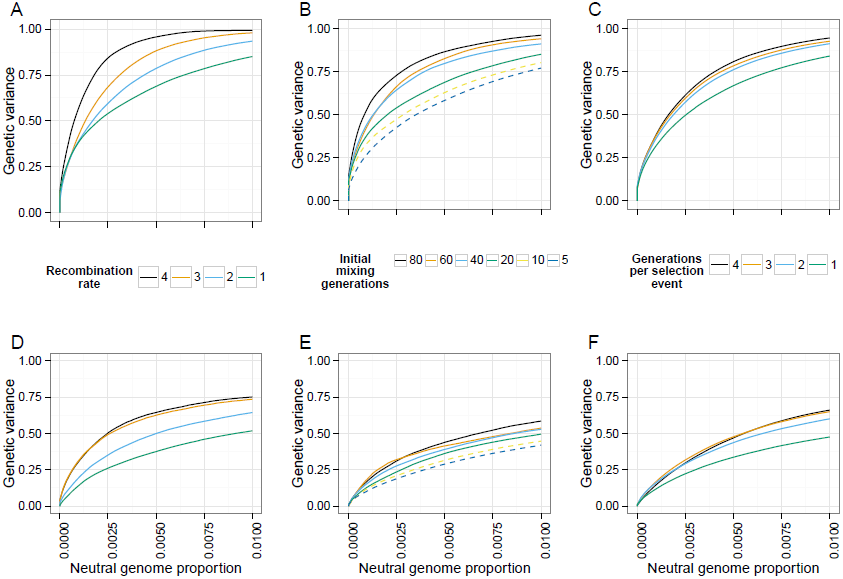
Effect of recombination. The power to detect and localize QTLs depends on the extent of recombination experienced by the populations. (A,D) Power increases with the recombination rate of the organisms under selection. (B,E) Additional generations of initial mixing and (C,F) additional generations between rounds of selection similarly increase power. [top row (A-C): 10 QTLs, bottom row (D-F): 100 QTLs, *h*^2^ = .5, N=1000, 20% selected, 20 generations of selection, 20 generations of mixing in A, C, D, F]

One way to decrease linkage disequilibrium in the population is to allow more generations of initial neutral mixing in the founder population, before selection starts (e.g. Turner and Miller (2012)). We simulated scenarios with varying numbers of generations of neutral mixing (20, 40, 60, 80), with the other parameters as above. We found that increasing the number of generations of initial neutral mixing increases power (Figure 8BE).

An alternative way to decrease linkage disequlibrium is to intersperse extra generations of neutral mixing between rounds of selection. To test this idea, we simulated scenarios where selection was carried out every 1, 2, 3, or 4 generations, with random mating during the generations where selection did not take place (other parameters as above). Again, we found that the additional recombination events increased power to detect QTLs (Figure 8CF). We further illustrate the effect of extra recombination on allele frequency differences in Figure 9.

**Figure 9.**
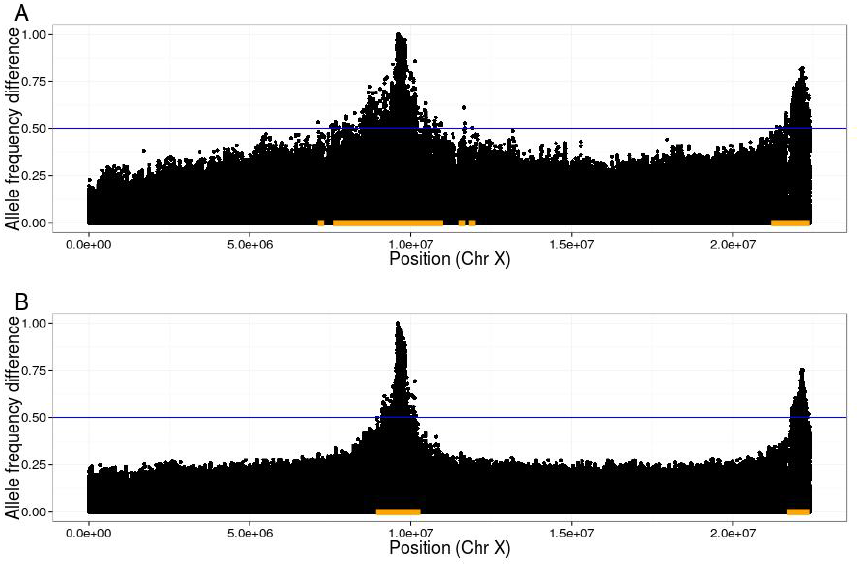
Illustration of linkage disequilibrium between QTLs and neutral loci. In plots of allele frequency differences between high and low populations, QTL peaks are narrower when extra generations of neutral mixing are introduced. As an example, we indicate a threshold of .5 with the blue lines, and the corresponding detection regions with orange bars. Note that the true region, consisting of sites within 10kb of either SNP, is too small to be seen at this scale; thus, the size of the orange bars represent the (local) false positive rate at this threshold. [10 QTLs, *h*^2^ = .5, N=1000, 20% selected, 20 generations of initial mixing, shown are average D values over 20 replicates. A) 20 generations of selection B) 20 generations of selection, 4 generations per selection event (80 generations total)]

Taken together, these results show that the ability to detect and localize QTLs depends crucially on recombination, both to decrease interference between QTLs and to break up linkage between QTLs and nearby neutral variants.

### Haplotype-based inference of allele frequencies increases power

In all of our power analyses, we have assumed that we are able to calculate the allele frequencies at all variant sites perfectly. However, in actual experiments, the allele frequency at a locus will be estimated by considering the sequence read counts that cover the site. These estimates are prone to errors from two sources: random base errors, and stochasticity in the number of reads covering the site. The error in measurement of allele frequencies will lead to a loss of power to detect and localize QTLs.

In the case where founder sequences are known, our previous work (Kessner *et al.*, 2013) showed that one can leverage haplotype information from the founders to obtain local haplotype frequencies, and that these estimates are robust to both sequence read errors and uneven coverage. This work suggested that improved allele frequency estimates could be derived from local haplotype frequency estimates.

We investigated the effect of allele frequency estimation error on power by first obtaining empirical error distributions for the two estimation methods (see Methods). We then simulated artificial selection experiments with our standard experimental parameters [*N* = 1000, 20% selected, 20 generations] and canonical trait architectures, followed by the random introduction of errors based on the empirical error distributions. In all cases, we found that the haplotype-based allele frequency estimates led to improved power over the read-count-based estimates. Figure 10A shows the results from the case with 10 QTLs and *h*^2^ = .5.

**Figure 10.**
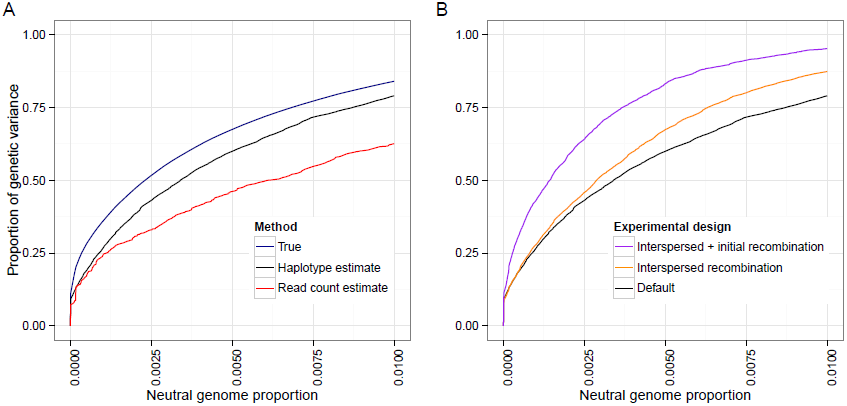
Using founder haplotype information estimation improves power. A) If founder sequence information is available, estimating local haplotype frequencies leads to better allele frequency estimates and an increase in power over estimates obtained from raw read counts. B) Error in local haplotype frequency estimates increases with the number of generations due to recombination; however, there is a still a net gain in power from extra generations of neutral mixing. Default: 20 generations mixing, 20 generations selection. Interspersed: 20 generations mixing, 4 generations per selection event, 20 selection events (100 generations total). Initial + interspersed: as interspersed, with 70 generations mixing (150 generations total). [10 QTLs, *h*^2^ = .5]

Our analyses from the previous section showed that increasing the amount of recombination led to an increase in power. On the other hand, increased recombination results in shorter haplotypes. This leads to greater errors in local haplotype frequency estimates and the allele frequency estimates derived from them, with a subsequent decrease in power. To investigate the relative magnitude of these two counteracting effects, we added two scenarios: one where we interspersed rounds of neutral mating between generations of selection (4 generations per selection event, 100 generations total for the experiment), and one where we also added an extra 50 generations of initial neutral mixing (150 generations total). We introduced random errors as above, using empirical error distributions that take into account the increased number of generations.

We found that recombination’s positive effect of breaking up linkage between QTLs and neutral sites outweighed the negative effect on haplotype frequency estimation (Figure 10B). However, for experiments lasting 200 generations or longer, our haplotype-derived allele frequency estimates were no better than read-count-derived estimates (see Methods for parameter assumptions). Thus, we do not expect haplotype-derived allele frequency estimates to lead to increased power in scenarios where the typical length of haplotype chunks is shorter than the window size used for the local haplotype frequency estimation.

## Discussion

We have presented a new analysis of the power of artificial selection experiments to detect and localize loci contributing to a quantitative trait. In this analysis, we explicitly model whole genomes of individuals, quantitative traits, and selection based on individual trait values, using a novel simulation framework.

We showed that population genetic simulations based on loci with constant selection coefficients do not fully capture the dynamics of QTLs contributing to a trait under artificial selection, and that the trait architecture plays a large role in these dynamics. In addition, explicit modeling of selection on a quantitative trait has several other advantages. For example, simulated experiments can be parameterized and results can be reported using the standard quantitative genetics concepts of effect size, genetic variance, and heritability. Also, scenarios such as divergent selection can be simulated in a straightforward manner. Finally, the behavior of a QTL under artificial selection is dependent both on the experimental design (proportion of individuals selected each generation) and on the trait architecture (effect size, and linkage to other QTLs). While the selection coefficient conflates these parameters into a single number, our simulation framework allows these parameters to be separated and investigated independently.

Our results show the important role that recombination plays in the ability to identify QTLs. Recombination not only reduces interference between QTLs, but also decreases linkage disequilibrium between QTLs and neighboring neutral loci. In fact, one can view the artificial selection experiment as a classification problem. From this viewpoint, the ROC curve, which measures the ability to classify a locus as QTL vs. neutral, contains information about how well the experiment can break linkage disequilibrium between causal and neutral loci.

The classification viewpoint is useful when thinking about how to design a selection experiment. For example, we found that experiments allowing more opportunity for recombination, either during initial neutral mixing of the founder haplotypes or between selection events, have greater power to detect and localize QTLs. Similarly, experiments with weaker selection (greater proportion selected each generation) over a longer period of time will have greater power than shorter experiments with strong selection. We note that the improved mapping resolution afforded by additional recombination is analagous to the use of recombinant inbred lines (see Crow (2007) for a history) or advanced intercross lines (Darvasi and Soller, 1995) in traditional QTL mapping studies.

While recombination increases the ideal power of an artificial selection experiment, it also decreases the ability to use founder haplotype information to obtain more accurate allele frequency estimates. We showed that the haplotype-based estimates still result in a net increase in power in experimental scenarios where the scale of recombination is larger than the window used for estimating local haplotype frequencies. This suggests that when choosing the window size for such an analysis, one should take into consideration the expected scale of recombination based on the experimental design.

Similar to the findings of Kofler and Schlotterer (2014) and Baldwin-Brown *et al.* (2014), we found that increasing population sizes and number of replicates leads to an increase in power. Additionally, our simulation framework allowed us to quantify the increase in power due to bidirectional selection. We also note that adding generations of initial neutral mixing in the founder population is in some ways similar to increasing the number of founder haplotypes, in that it places QTLs on multiple genetic backgrounds. Our results regarding the increase in power due to additional initial mixing are thus consistent with the findings of both of these groups that increasing the number of founder haplotypes increases power to detect and localize QTLs.

Also in agreement with Baldwin-Brown *et al.* (2014), but in contrast to Kofler and Schlotterer (2014), we found that the effect of increasing the length of the experiment is not uniformly beneficial, but rather depends on the strength of selection. From the classification viewpoint again, power depends on the ability of the experiment to allow QTLs to differentiate in selected populations while keeping allele frequencies at neutral loci constant. Continuing the experiment beyond the point where the majority of QTLs have differentiated will lead to increased fixation of linked neutral loci, and hence lower power.

In contrast to both Kofler and Schlotterer (2014) and Baldwin-Brown *et al.* (2014), whose results suggest that hundreds of generations are necessary to obtain reasonable power, we found that artificial selection experiments can detect QTLs explaining most of the genetic variance of the trait in as little as 20 generations, under reasonable assumptions about the trait architecture (100 loci, *h*^2^ = .5, exponentially distributed effect sizes). We believe that the previous studies were perhaps conservative in their choice of ranges of selection coefficients. On the other hand, it is not obvious how to interpret selection coefficients in the context of an artificial selection on a quantitative trait, where we feel it is more natural to use parameters representing the trait architecture and experimental design.

In summary, we have shown that it is feasible to do whole-genome simulations of artificial selection with explicit quantitative trait modeling. The scope of this study was limited to the use of *D. melanogaster* as our model species and simple trait architectures with no dominance or pleiotropic effects. However, we believe that our simulation methodology can be applied to a wide variety of species, trait architectures, and experimental protocols. We emphasize that the opportunity for recombination is a key factor in the power to detect and localize QTLs, and that this should be taken into account by future designers of artificial selection experiments.

## Methods

### Forward simulation

We used the program forqs (Kessner and Novembre, 2014) for all forward simulations. forqs simulates whole genomes of individuals efficiently by tracking the haplotype chunks that are inherited from the founder individuals in the initial generation. forqs allows the specification of quantitative traits, and provides flexible fitness functions for the simulation of complex evolutionary scenarios such as artificial selection.

In our simulations of artificial selection on a quantitative trait, individuals had 3 chromosomes, with lengths matching *Drosophila* chromosomes X, 2, and 3. For simplicity, we set the recombination rate to be uniform along each chromosome, with rate = 2 cM/mb, which is similar to recombination rates reported previously (Comeron *et al.*, 2012). For each set of experimental parameters we simulated populations using 200 replicates each of 12 different canonical architectures: 2, 5, 10, and 100 QTLs, at initial heritability levels .2, .5, and .8. For a given number of QTLs and heritability level, QTL positions and effect sizes were generated randomly for each simulation run (see next section). The random trait generation was implemented in an auxilliary program which produces trait description files which are included by forqs configuration files that specify the experimental setup. Our simulation scenarios represented selection on standing variation where the expected waiting time 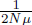 for a mutation to occur at a QTL is much larger than the number of generations in the experiment. Hence, we did not include *de novo* mutations in our simulations.

Each generation, forqs calculates trait values for all individuals based on the effect sizes of the alleles they carry at QTLs and a random environmental effect (with variance determined by the heritability of the trait). Artificial selection was simulated using the forqs module FitnessFunction TruncationSelection, which assigns a fitness value of 1 to those individuals whose trait value lies above (or below) a threshold (and 0 otherwise), where the threshold is determined each generation by the distribution of trait values in the population and the user-specified proportion of individuals selected to create the next generation.

All configuration files and analysis scripts for all simulations are freely available online: https://bitbucket.org/dkessner/artificial_selection_pipeline

### Generation of random trait architectures

In order to investigate how the genetic architecture of a trait affects the behavior of QTL allele frequencies under artificial selection, we developed a method to generate random trait architectures with specified parameters. In particular, we were interested in how the number of QTLs contributing to the trait and the heritability of the trait affect the power to detect QTLs. In addition, we wanted to investigate how the trait architecture affects a focal QTL with a specified effect size and initial allele frequency. Finally, we wanted to ensure that linkage disequilibrium in the simulated starting population is similar to that found in populations used in experimental settings. In the following, we describe our procedure for generating the founding population and trait architecture in our simulations.

Starting with founder individuals for which we have full-genome haplotypes, we run neutral forward simulations with forqs to recombine the haplotypes and expand the population size. This results in a mixed population that is 2-3x larger than the desired starting population for the selection experiment. For our founder individuals, we used the publicly available SNP data from 162 Drosophila inbred lines representing Freeze 1 of the Drosophila Genetic Reference Panel (DGRP) project (Mackay *et al.*, 2012). Our procedure is similar to the experimental procedure used by Turner and Miller (2012), where individuals from the DGRP inbred lines are allowed to mate randomly for several generations in order to create a mixed population with genetic variation and linkage disequilibrium similar to the natural popluation from which the inbred lines were derived.

In some cases we additionally specify a focal QTL, for which we specify the locus, effect size and initial allele frequency. We create a starting population by randomly selecting a subset of individuals from the mixed population. We ensure that the focal QTL has the desired allele frequency by choosing individuals in Hardy-Weinberg proportions according to their focal QTL genotype.

From the heritability and total variance parameters that we specify, we calculate a target genetic variance for the trait. We choose the remaining QTL positions uniformly at random across the genome, until we have the specified number of QTLs. We choose effect sizes according to a standard exponential distribution, with random sign for positive/negative effect on the trait. From the individuals’ haplotypes and QTL effect sizes, we calculate the current genetic variance of the trait in the population. We then scale the QTL effect sizes so that the genetic variance is equal to the target genetic variance. In the case where we have a focal QTL, we keep the focal QTL effect size constant, and scale the other QTL effect sizes – this requires iteration until the genetic variance is close to the target genetic variance, within a specified tolerance.

### Realized selection coefficient

We first recall the deterministic model of selection on a single locus in a diploid population. Suppose a locus has two alleles *A*_0_ and *A*_1_. If *p* is the frequency of *A*_1_, then the allele frequency change is given by:

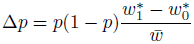

where *w*_0_^*^ and *w*_1_^*^ are the marginal fitnesses of the *A*_0_ and *A*_1_ alleles, respectively, and 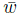 is the mean fitness of the population (Rice, 2004).

In the special case where *A*_1_ has additive selection coefficient *s*, this becomes:

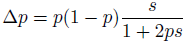

We define the *realized selection coefficient* for a given *p* and Δ*p* to be the selection coefficient *s* that would result in this allele frequency change in the deterministic case. By solving the above equation for *s*, we obtain a formula for the realized selection coefficient:

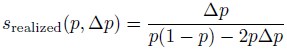

### Calculation of power and false positive rate

We explored several methods for calculating the power and false positive rate associated with the detection of QTLs by allele frequency differences between populations.

The simplest method to measure power is to calculate the proportion of QTLs detected. However, we feel that a more relevant measure of power is the proportion of the genetic variance in the initial population that is explained by the detected QTLs. This measure appropriately gives greater weight to QTLs responsible for more of the genetic variance.

Figure 11AB illustrates the difference between these two measures of power: at a false positive rate of 10*^−^*^4^, only 50% of the QTLs are detected, but these QTLs are responsible for more than 75% of the genetic variance of the trait.

**Figure 11.**
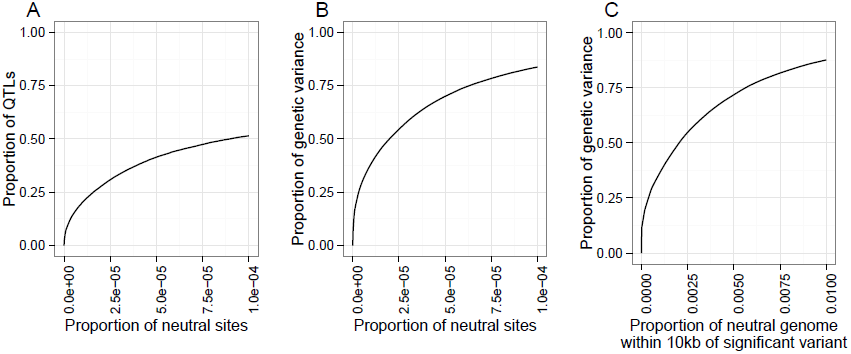
Comparison of three methods for calculating and interpreting power and false positive rate. A) Power is measured by proportion of QTLs detected, and false positive rate is measured by proportion of neutral variant sites detected. B) Power is measured by the proportion of genetic variance in the founder population explained by the detected QTLs; false positive rate as in A. C) Power as in B; false positive rate is measured by proportion of the neutral genome covered by the detection region. In this case, the ROC curve shows that over 75% of the genetic variance is explained by QTLs in a detection region which covers 1% of the neutral genome.

Furthermore, calculating the false positive rate as the proportion of neutral variants detected does not capture the clustering of detected variants due to linkage disequilibrium. In addition, significantly diverged variants are not necessarily causal, but indicate that the region nearby may contain a causal variant. To address these issues, we define a detection region to include all variants within a specified radius of any variant whose *D* value exceeds a given threshold (see Results, “Measurement of power to detect and localize QTLs”). This leads to a natural definition of false positive rate: the proportion of the neutral genome covered by the detection region (see Figure 9 for an illustration).

We calculated the false positive rate in this way with radii of 10kb, 100kb, and 1mb. We found that using a radius of 100kb or 1mb led to true regions that covered a substantial portion of the genome in the cases where there were a large number of QTLs. Thus, we found that using the 10kb radius for the detection region, together with measuring power as the proportion of variance explained, gave the most interpretable results.

### Analysis pipeline

The forward simulator forqs efficiently simulates the entire genome of each individual by tracking haplotype chunks. forqs outputs data files that represent each chromosome of each individual as a mosaic of founder haplotypes. In order to obtain the neutral variants carried by an individual, the neutral variation on founder chromosomes must be propagated to the individual’s mosaic chromosomes.

In our simulation framework, we use two forward simulations. The first simulation creates a mixed population from the founder haplotypes, after which we generate the random trait architecture. The second simulation represents the selection experiment. Individuals in the final populations are mosaics of individuals in the mixed population, which are in turn mosaics of the founders. We implemented a custom program to handle this two-step propagation of neutral variation. Given founder sequences, the mixed population, and the final population, the program calculates the allele frequency in the final population of each variant in the genome. We note that while this step does not require a large amount of computation, it is I/O-intensive because it uses the full DGRP variant data set for the founder sequences, which consists of 162 haplotypes at approximately 5 million loci.

After calculating allele frequencies for each population, we calculate the allele frequency difference *D* for each high-low population pair under consideration (multiple pairs for the replication analyses). After sorting the *D* values, we begin with the highest *D* value and iteratively decrease the threshold to obtain data equivalent to an ROC curve (power and false positive rates) for that simulation run. We note that this step depends on the method for calculating power and false positive rate, so we performed it once for each method we described above in “Calculation of power and false positive rate”.

We obtain average ROC curves by calculating the average power over replicate simulation runs at regularly spaced false positive rates, where the power for a particular run at a given false positive rate is obtained by linear interpolation between points on its ROC curve.

### Empirical error distributions

To investigate the power increase due to the use of haplotype-based allele frequency estimates, we needed to simulate pooled sequence reads from a population, followed by haplotype frequency estimation in sliding windows across the genome, using the harp method and software detailed in Kessner *et al.* (2013).

Because this procedure is computationally expensive, and due to the large number of simulations involved in this study, it was not feasible to do this for each simulated experiment.

As an alternative, we obtained empirical error distributions, which we later used to add random errors to true allele frequencies. Because errors in the haplotype frequency estimation depend on the length scale of recombination, we ran replicate neutral simulations for varying numbers of generations (from 40 to 400). Haplotypes surrounding selected QTLs are expected to be longer than in neutral regions, so our empirical error distributions are conservative. We simulated pooled sequence reads from the final populations using a new program simreads forqs based on the read simulator used in Kessner *et al.* (2013). Haplotype frequency estimation was performed with harp in overlapping sliding 200kb windows within a single 1mb region. We then calculated allele frequencies at variant sites within the region using read counts. By considering allele frequencies in bins of size .05, we obtained a frequency-dependent empirical error distribution. Similarly, we also derived allele frequency estimates from the local haplotype frequencies, from which we obtained frequency-dependent empirical error distributions for each generation count. We found that after ∼ 200 generations, the haplotype-based estimates were no better than the read-count-based estimates; this behavior is expected as the recombination length scale approaches the window size used for haplotype frequency estimation (200kb in this analysis). Hence, for this analysis we only considered experimental scenarios lasting less than 200 generations.

## Acknowledgments

The authors would like to thank Alex Platt, Charleston Chiang, Eunjung Han, and Diego Ortega Del Vecchyo for helpful comments and discussion.

